# Functional heterogeneity and plasticity in naïve CD8 T cells drive superior effector and memory responses

**DOI:** 10.1101/2025.10.16.682909

**Authors:** Yamato Sajiki, Satomi Ando, Charles M. Perkins, Yi-Chung Huang, Katie E. Smith, Carolyn M. Doerning, Koichi Araki

## Abstract

While significant progress has been made in defining subsets among antigen-experienced CD8 T cells, the heterogeneity of naïve CD8 T cells remains poorly understood. Here, we identify naïve CD8 T cell subsets with superior persistence and an enhanced capacity to generate effector and memory cells, leading to more effective protection. These high-quality naïve CD8 T cells are marked by IL-18Rα, CD73, and CXCR3, and functionally less potent naïve CD8 T cells can convert into these superior subsets. Their enhanced response to infections is driven by better survival of the progeny effector cells during the T cell expansion phase. This improved survival is mediated by increased Ly6C2 expression on effector cells derived from these high-quality naïve cells. Collectively, our findings reveal functional heterogeneity and plasticity among naïve CD8 T cells and uncover a mechanism by which high-quality naïve subsets drive robust CD8 T cell responses, providing a previously unrecognized layer of immune regulation.

**SUMMARY:** Our study reveals functional heterogeneity and plasticity within naïve CD8 T cells, identifying high-quality subsets that can emerge from less potent cells and drive superior immune responses through enhanced survival of effector progeny.

## INTRODUCTION

CD8 T cells play an essential role in controlling viral, intracellular bacterial, and parasitic infections as well as tumors (Appay et al., 2008; Laidlaw et al., 2016; Padilla et al., 2009; St Paul and Ohashi, 2020). Over the past few decades, considerable efforts have been devoted to defining subpopulations of activated CD8 T cells, leading to the identification of multiple subsets within effector, memory, and exhausted CD8 T cells (Hashimoto et al., 2018; Kaech and Cui, 2012; Martin and Badovinac, 2018). These findings advanced our understanding of CD8 T cell heterogeneity in acute and chronic infections and cancer by clarifying the distinct roles of each subset (Chen et al., 2018; Hashimoto et al., 2018; Kaech and Cui, 2012; Martin and Badovinac, 2018; McLane et al., 2019). Additionally, these studies have revealed that phenotypically distinct subsets exhibit differential functions such as cytokine production, proliferative capacity, longevity, and multipotency. Thus, dissecting the heterogeneity of CD8 T cells is crucial for understanding their responses to antigen-stimulation and mechanisms of maintenance.

In contrast to antigen-experienced T cell populations, studies of naïve T cells have generally focused on quantity over quality (den Braber et al., 2012; Zlamy et al., 2016), as they were traditionally considered a homogeneous population. However, recent studies have revealed functional heterogeneity within the naïve CD4 T cell population (Deep et al., 2024; ElTanbouly et al., 2020; Even et al., 2024; Rogers et al., 2021; Yoon et al., 2025), suggesting that naïve T cells could be viable targets for enhancing T cell immunity. For example, if a functionally inferior naïve T cell population is primarily activated at the time of vaccination, vaccine-induced immunity is likely to be suboptimal. Furthermore, if naïve T cell subsets with impaired ability to generate a sufficient number of effector cells can be transformed into normal or functionally superior naïve T cells, the overall T cell immune response can be improved. Since naïve T cells have an enormous TCR repertoire diversity capable of recognizing virtually any non-self antigens including pathogens and tumor neoantigens (de Greef et al., 2020; Qi et al., 2014), creating a high-quality naïve CD8 T cell population ready to give rise to potent long-lasting T cell responses would provide better protection against infections, enhance vaccine-induced immunity, and prevent cancer. Achieving these goals requires defining and gaining a deeper understanding of naïve CD8 T cell heterogeneity.

Despite several studies examining this topic (Berton et al., 2023; Eggert et al., 2024; Fulton et al., 2015; Jergovic et al., 2021; Ju et al., 2021; Lynch et al., 2025), our understanding of naïve CD8 T cell heterogeneity remains limited compared to that of memory CD8 T cells. It is well established that effector memory CD8 T cells can convert to central memory cells with superior longevity and protective capacity due to their enhanced proliferative potential (Wherry et al., 2003). Whether a similar differentiation process occurs within the naïve CD8 T cell pool remains unclear. To address this, we adoptively transferred naïve CD8 T cells into recipient mice and observed dynamic changes in the quality of antigen-specific naïve CD8 T cells over time.

Initially, only a small fraction of freshly isolated naïve CD8 T cells exhibited an IL- 18Rα^Hi^CXCR3^Hi^CD73^Hi^ (triple-positive) phenotype. However, the frequency of triple-positive naïve CD8 T cells increased substantially one month after adoptive transfer into recipient mice without any infection or stimulation. These long-lived triple-positive naïve CD8 T cells demonstrated an extended lifespan compared to the freshly isolated naïve CD8 T cells and produced greater numbers of both effector and memory cells upon infection. Their increased response to infections was driven by better survival of effector cells during the T cell expansion phase, and we identified Ly6C2 as a key regulator of this process. Collectively, our findings demonstrate functional heterogeneity within the naïve CD8 T cell population and provide a foundation for developing novel approaches to enhance CD8 T cell immunity by targeting the naïve T cell population.

## RESULTS

### Naïve CD8 T cells undergo functional changes over time

Effector memory CD8 T cells are known to convert to central memory cells over time, as demonstrated by adoptive transfer experiments into recipient mice (Wherry et al., 2003). To investigate whether naïve CD8 T cells similarly undergo functional and phenotypic changes over time, we adoptively transferred P14 cells, which are T cell receptor (TCR) transgenic CD8 T cells specific for the lymphocytic choriomeningitis virus (LCMV) GP33 epitope (Pircher et al., 1989), in the absence of infection or cognate antigen stimulation (Fig. 1A). We monitored the survival kinetics of the transferred P14 cells and observed that their frequency reduced to 50% within the first month. However, it required over 2 additional months for the remaining P14 cells to decrease from 50% to 25% (Fig. 1B), indicating improved survival of P14 cells after the initial one month post-transfer. To examine whether this improved survival is associated with phenotypic changes, we analyzed several naïve CD8 T cell markers on P14 cells before transfer and at 35 days post-transfer. The expression of CD44, a marker that distinguishes memory from naïve CD8 T cells (Baaten et al., 2012), remained low on P14 cells post-transfer compared to infection-induced LCMV-specific memory CD8 T cells (Fig. 1C), indicating that P14 cells housed in B6 mice retained a naïve phenotype. While CD62L expression, a lymphoid tissue homing receptor, remained unchanged (Fig. 1C), CD127 expression, the IL-7 receptor that is essential for naïve T cell survival (Takada and Jameson, 2009), was modestly upregulated on P14 cells 35 days after transfer (Fig. 1C), potentially contributing to the improved survival seen beyond the first month post-transfer. These observations gave rise to the hypothesis that naïve P14 CD8 T cells that exhibit survival advantage may also show discordant response to antigenic stimulation. To address this, P14 cells housed in B6 mice for over 30 days (hereafter referred to as long-lived P14 cells) were re-isolated and co-transferred with an equal number of freshly isolated P14 cells into new recipient B6 mice, followed by LCMV-Armstrong (LCMV-Arm) infection, which causes an acute infection (Fig. 1D). Compared to fresh P14 cells, long-lived P14 cells produced significantly more effector cells on day 8 post-infection (Fig. 1, E and F).

**Fig. 1.**
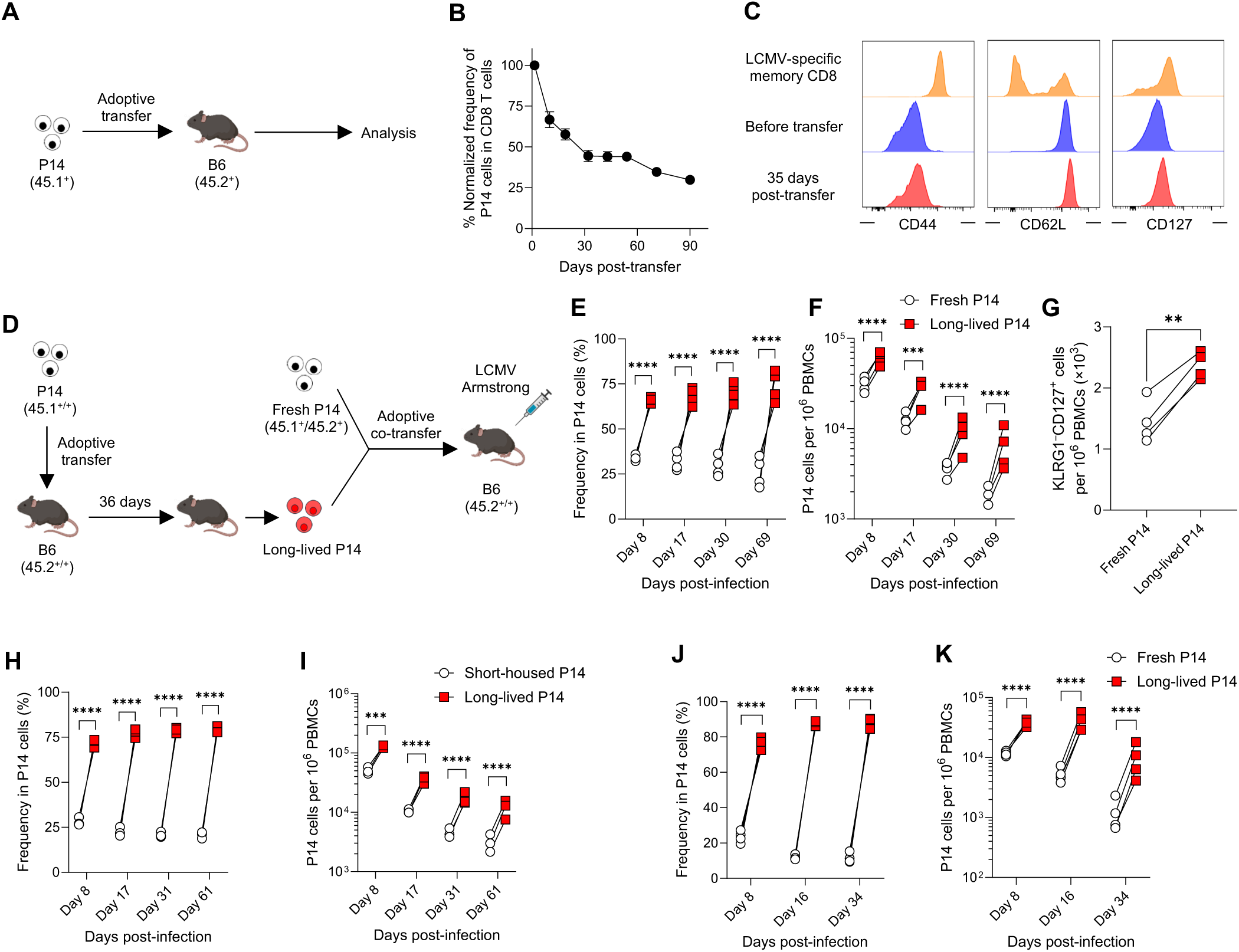
Long-lived naïve CD8 T cells exhibit superior responses following infection. (A) Experimental design for panel B. P14 CD8 T cells (5 × 10^6^ cells, CD45.1^+^) were adoptively transferred into B6 mice (CD45.2^+^), and their kinetics were monitored in the blood. (B) Kinetics of adoptively transferred P14 CD8 T cells (CD45.1^+^ DbGP33 tetramer^+^ CD8 T cells) in the blood without infection. The frequency of P14 CD8 T cells at 1.5 days post-transfer was set as 100%. Each symbol represents the mean and error bars indicate SEM. (C) CD44, CD62L, and CD127 expression on splenic P14 CD8 T cells before transfer and at 35 days post-transfer. As a control, the expression of these markers on DbGP33^+^ memory CD8 T cells obtained from the spleen of LCMV-Arm infected mice (> day 100 post-infection) is shown. Histograms were gated on DbGP33^+^CD8 T cells. (D) Experimental design for panels E to G. P14 CD8 T cells (long-lived P14, CD45.1^+/+^), housed for 36 days after transfer in B6 mice, were mixed with an equal number of freshly isolated P14 CD8 T cells (fresh P14, CD45.1^+^CD45.2^+^). The mixture (1 × 10^3^ P14 CD8 T cells of each population) was adoptively transferred into CD45.2^+/+^ B6 mice, followed by LCMV-Arm infection. (E and F) Percentages of progeny derived from fresh and long-lived P14 CD8 T cells within total P14 CD8 T cells (E) and their kinetics (F) in the blood post-infection. (G) The number of memory precursor effector P14 cells (CD127^+^KLRG1^−^) in the blood on day 8 post-infection. (H and I) Equal numbers (1 × 10^3^ cells each) of congenically distinct P14 CD8 T cells, housed for 1.5 days (short-housed P14) and 35 days (long-lived P14), were adoptively co-transferred into B6 mice, followed by LCMV-Arm infection. Percentages of progeny derived from each population within total P14 CD8 T cells (H) and their kinetics (I) in the blood post-infection. (J and K) Responses of long-lived P14 cells during chronic LCMV-Clone 13 infection. Long-lived P14 CD8 T cells (1 × 10^3^ cells, CD45.1^+/+^), housed for 74 days in B6 mice, were adoptively co-transferred with an equal number of fresh P14 CD8 T cells (1 × 10^3^ cells, CD45.1^+^CD45.2^+^) into B6 mice (CD45.2^+/+^), followed by LCMV-Clone 13 infection. Percentages of progeny derived from each population within total P14 CD8 T cells (J) and their kinetics (K) in the blood post-infection. Data are representative of 2 or more independent experiments with 3 or more mice per group. In E–K, each symbol represents an individual mouse, and lines indicate paired comparisons within the same mice. Statistical analysis was performed using paired *t* test (G) and one-way ANOVA (E, F, H, I, J, and K). ** *p* < 0.01; *** *p* < 0.001; **** *p* < 0.0001.

Although the proportions of memory precursor effector cells (CD127^+^KLRG1^−^) and short-lived effector cells (CD127^−^KLRG1^+^) (Huster et al., 2004; Joshi et al., 2007; Kaech et al., 2003; Sarkar et al., 2008) were similar between the two groups (Fig. S1A), the number of memory precursor effector cells derived from long-lived P14 cells was significantly higher (Fig. 1G).

Consequently, long-lived P14 cells gave rise to a significantly larger memory T cell pool compared to fresh P14 cells (Fig. 1, E and F, Fig. S1, B and C). To rule out the possibility that adoptive transfer itself changes the function of P14 cells, long-lived P14 cells were co-transferred with an equal number of short-housed P14 cells (1.5 days post-transfer) into naïve B6 mice, followed by LCMV-Arm infection. Long-lived P14 cells showed an enhanced response compared to short-housed P14 cells (Fig. 1, H and I), demonstrating that a certain period after adoptive transfer is required to acquire the improved expansion capacity. Next, to determine if our findings from the acute viral infection model with LCMV-Arm could be generalized to other systems, we examined the response of long-lived P14 cells to chronic viral infection with LCMV-Clone 13 and acute bacterial infection with GP33 epitope-expressing *Listeria monocytogenes*. Consistent with LCMV-Arm infection, long-lived P14 cells exhibited substantial expansion compared to fresh P14 cells during both LCMV-Clone 13 and *Listeria monocytogenes* infections (Fig. 1, J and K, Fig. S2, A–C). Taken together, these findings suggest that the function of antigen-specific naïve CD8 T cells changes over time, contributing to their heterogeneity.

### Naïve CD8 T cells change their phenotype over time

Previous studies have identified markers such as CD5 and Ly6C that are associated with superior naïve CD8 T cell responses upon antigen stimulation (Berton et al., 2023; Fulton et al., 2015; Ju et al., 2021). To explore this, we compared CD5 and Ly6C expression between freshly isolated and long-lived P14 cells. While CD5 expression was indistinguishable between the two populations (Fig. S3A), Ly6C expression was upregulated in long-lived P14 cells compared to fresh P14 cells (Fig. S3A). To test whether the presence of the Ly6C^Hi^ subset in long-lived P14 cells accounts for their superior response upon infection, we sorted Ly6C^Lo^ and Ly6C^Hi^ long- lived P14 cells and co-transferred them with an equal number of fresh P14 cells into recipient mice, followed by LCMV-Arm infection (Fig. S3B). Both Ly6C^Lo^ and Ly6C^Hi^ long-lived P14 cells exhibited superior responses to LCMV infection compared to fresh P14 cells (Fig. S3C), indicating that the enhanced response of long-lived naïve P14 cells cannot be attributed solely to Ly6C expression status. Thus, to further characterize long-lived naïve P14 CD8 T cells prior to infection, in comparison to fresh naïve P14 CD8 T cells, we performed RNA sequencing (RNA-seq) analysis (Fig. 2A). Sample-to-sample distance analysis revealed distinct transcriptional profiles, with the two populations forming separate clusters (Fig. 2B). Using a fold-change cutoff of 1.6 and an adjusted *p*-value threshold of 0.01, we identified 90 upregulated and 79 downregulated genes in long-lived P14 cells compared to fresh P14 cells (Fig. 2C, Supplemental Table 1). Notably, genes associated with immune responses, such as *Il18r1* (Kaplanski, 2018), *Cxcr3* (Groom and Luster, 2011), and *Nt5e* (Bono et al., 2015), exhibited significant differences (Fig. 2, C–F). To validate these findings, we assessed protein expression levels of these molecules on splenic fresh and long-lived P14 cells. Flow cytometric analysis showed higher expression of IL-18 receptor alpha (IL-18Rα, encoded by *Il18r1*), CXCR3, and CD73 (encoded by *Nt5e*) in long-lived P14 cells compared to fresh P14 cells (Fig. 2, G–J). This phenotypic trend was consistent across long-lived P14 cells isolated from other tissues including livers, lymph nodes (LNs), and blood (Fig. S4). Based on differential expression of these three molecules, we were able to classify fresh P14 cells into several subsets (Fig. 2K). While most of the IL-18Rα^Lo^ P14 cells lacked CXCR3 and CD73 expression, 4 distinct subsets were identified among the IL- 18Rα^Hi^ P14 cells based on the expression levels of CXCR3 and CD73 (Fig. 2K).

**Fig. 2.**
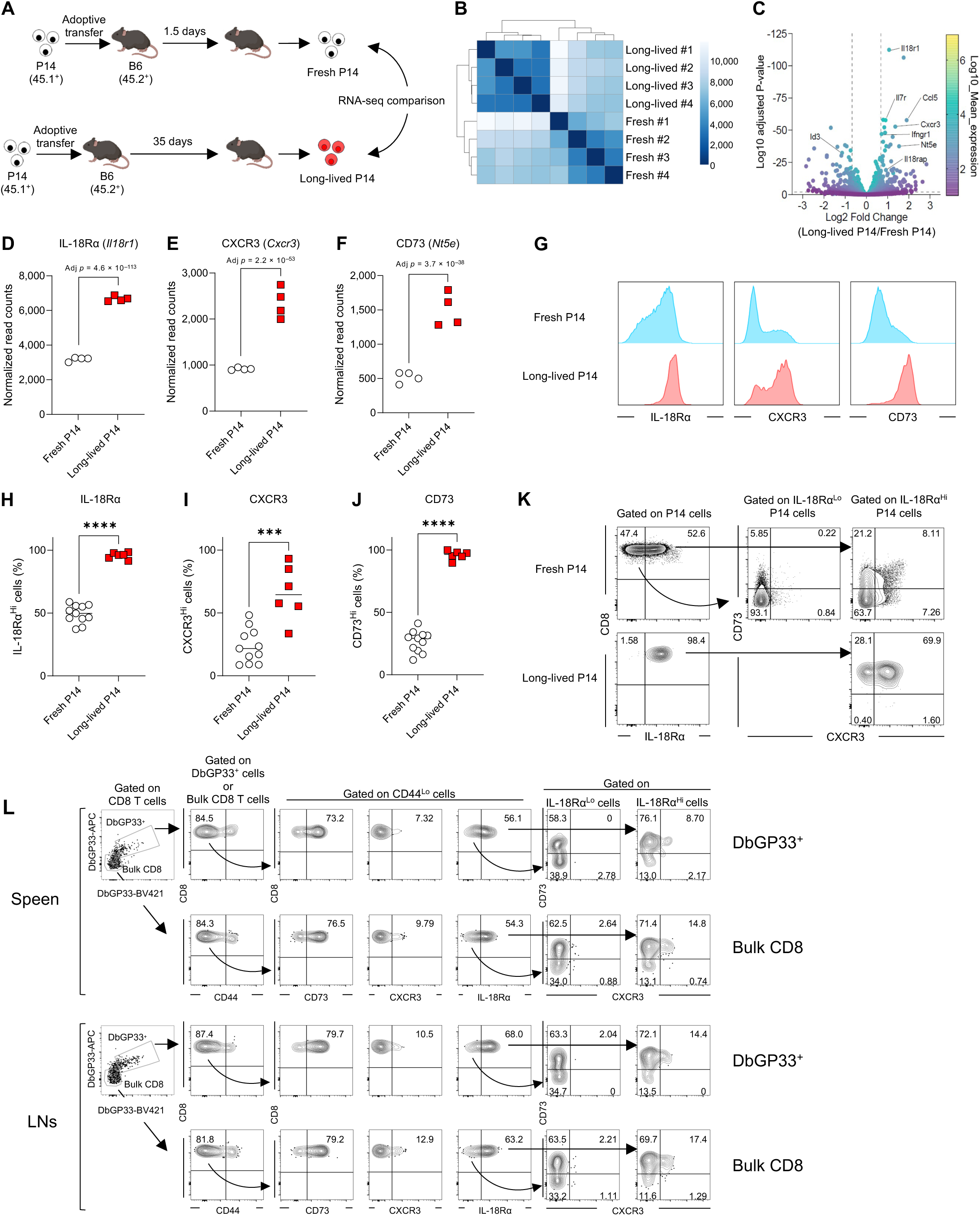
Long-lived naïve CD8 T cells are transcriptionally and phenotypically distinct from fresh naïve CD8 T cells. (A) Experimental design for panels B–F. P14 CD8 T cells (CD45.1^+^) were adoptively transferred into B6 mice (CD45.2^+^). On 1.5 days (fresh) and 35 days (long-lived) post-transfer, P14 CD8 T cells (CD45.1^+^DbGP33^+^CD8 T cells) were sorted from the spleens of recipient mice (*n* = 4 each) for RNA-seq analysis. Gene expression (read counts) was analyzed using DESeq2. (B) Heatmap of sample-to-sample distances using the Poisson Distance. (C) Fold changes and adjusted *p*-values of gene expression in long-lived P14 CD8 T cells relative to fresh P14 CD8 T cells. (D–F) Normalized read counts of *Il18r1* (D), *Cxcr3* (E), and *Nt5e* (F) in long-lived and fresh P14 CD8 T cells. Each symbol represents an individual mouse. (G–K) Protein expression levels of IL-18Rα, CXCR3, and CD73 on long-lived (>30 days post-transfer) and fresh P14 CD8 T cells in the spleens. (G) Representative histograms were gated on DbGP33^+^ P14 CD8 T cells. (H to J) Data were pooled from 6 independent experiments, and each symbol represents an individual mouse. (K) Heterogeneity of naïve P14 CD8 T cells (gated on DbGP33^+^ P14 CD8 T cells) based on the expression of IL-18Rα, CXCR3, and CD73. (L) Heterogeneity of endogenous bulk and DbGP33^+^ naïve CD8 T cells. Tetramer enrichment was performed using the cells isolated from the spleen and LNs (inguinal and brachial LNs) of naïve B6 mice, and the expression of IL-18Rα, CXCR3, and CD73 on either DbGP33^+^ or bulk CD44^Lo^CD8 T cells was analyzed. Statistical analyses were performed using Walt’s test (D–F) and unpaired t test (H–J). *** *p* < 0.001; **** *p* < 0.0001.

To determine whether similar heterogeneity exists in non-transgenic naïve CD8 T cells, we analyzed bulk CD44^Lo^ CD8 T cells in B6 mice and identified comparable subsets (Fig. 2L). However, since bulk CD44^Lo^ populations may include antigen-experienced cells – for example, exhausted CD8 T cells are known to downregulate CD44 – we further examined endogenous LCMV GP33-specific naïve CD8 T cells by tetramer enrichment (Moon et al., 2007; Obar et al., 2008). This analysis revealed that the multiple subsets observed in transgenic naïve CD8 T cells also exist among endogenous tetramer^+^ naïve CD8 T cells in spleens and LNs of B6 mice (Fig. 2L).

Next, to investigate whether heterogeneity of non-transgenic naïve CD8 T cells changes over time as observed in the P14 TCR transgenic CD8 T cells, we performed parabiosis, enabling effective naïve T cell transfer. By separating the fused mice 5 weeks after parabiosis, migration of naïve T cells from the other mouse was stopped and we were able to assess phenotypic changes of transferred naïve CD8 T cells (Fig. 3A). We found that IL-18Rα, CD73, and CXCR3 expression on transferred CD44^Lo^ CD8 T cells (both bulk and tetramer^+^ cells) increased compared to host counterparts (Fig. 3, B and C). These findings, including data from both transgenic and endogenous CD8 T cell experiments, demonstrate that naïve CD8 T cells exhibit phenotypic heterogeneity which can be altered over time.

**Fig. 3.**
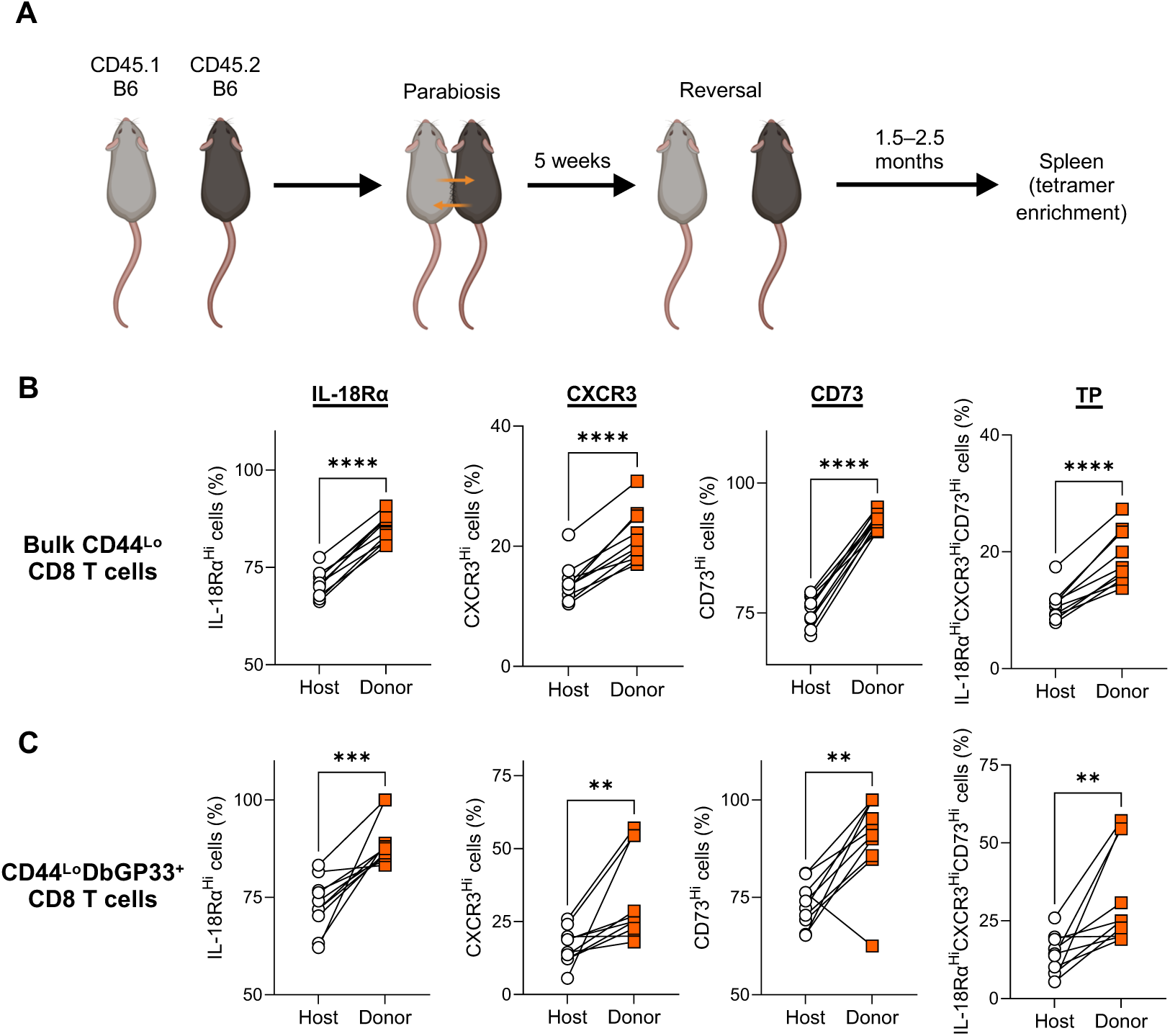
Endogenous naïve CD8 T cells change their phenotype over time. (A) Experimental design for panels B and C. Congenically distinct B6 mice were conjoined by parabiosis to effectively transfer donor CD8 T cells into another mouse. Parabiotic mice were separated 5 weeks after conjoining, which allowed for the prolonged monitoring of donor-derived CD8 T cell phenotype. 1.5 to 2.5 months after fusion reversal, the phenotype of donor- and host-derived bulk CD44^Lo^ (B) and DbGP33^+^CD44^Lo^ (C) CD8 T cells in the spleen was analyzed using tetramer enrichment. Data are pooled from 3 independent experiments with 10 mice. Each symbol represents an individual mouse and lines indicate paired comparisons within the same mouse. Statistical analysis was performed using paired *t* test. ** *p* < 0.01; *** *p* < 0.001; **** *p* < 0.0001.

### Plasticity and lineage relationship in naïve CD8 T cell subsets

In contrast to fresh P14 cells, long-lived P14 cells exhibited high frequency of a triple positive (TP, IL-18Rα^Hi^CXCR3^Hi^CD73^Hi^) subset (Fig. 2K). This raised the question of how long-lived P14 cells acquired this phenotypic feature. One possibility is that a triple negative (TN, IL- 18Rα^Lo^CXCR3^Lo^CD73^Lo^) population differentiates into the TP CD8 T cells over time. To address this question, TP, TN, and unsorted fresh P14 cells were separately transferred into B6 mice and the phenotype of each P14 population was monitored (Fig. 4A). While TP P14 cells maintained the expression levels of these molecules throughout the observation period, the TN population gradually upregulated them over time (Fig. 4B). Importantly, this differentiation mostly occurred in P14 cells that had not yet divided (Fig. 4C), indicating that the emergence of triple positive cells from the triple negative population truly represents a conversion of the two subsets rather than the proliferation of a few contaminating triple positive cells within the purified triple negative population. These results suggest that naïve CD8 T cells acquire the TP phenotype through differentiation, establishing a lineage relationship between TN and TP populations.

**Fig. 4.**
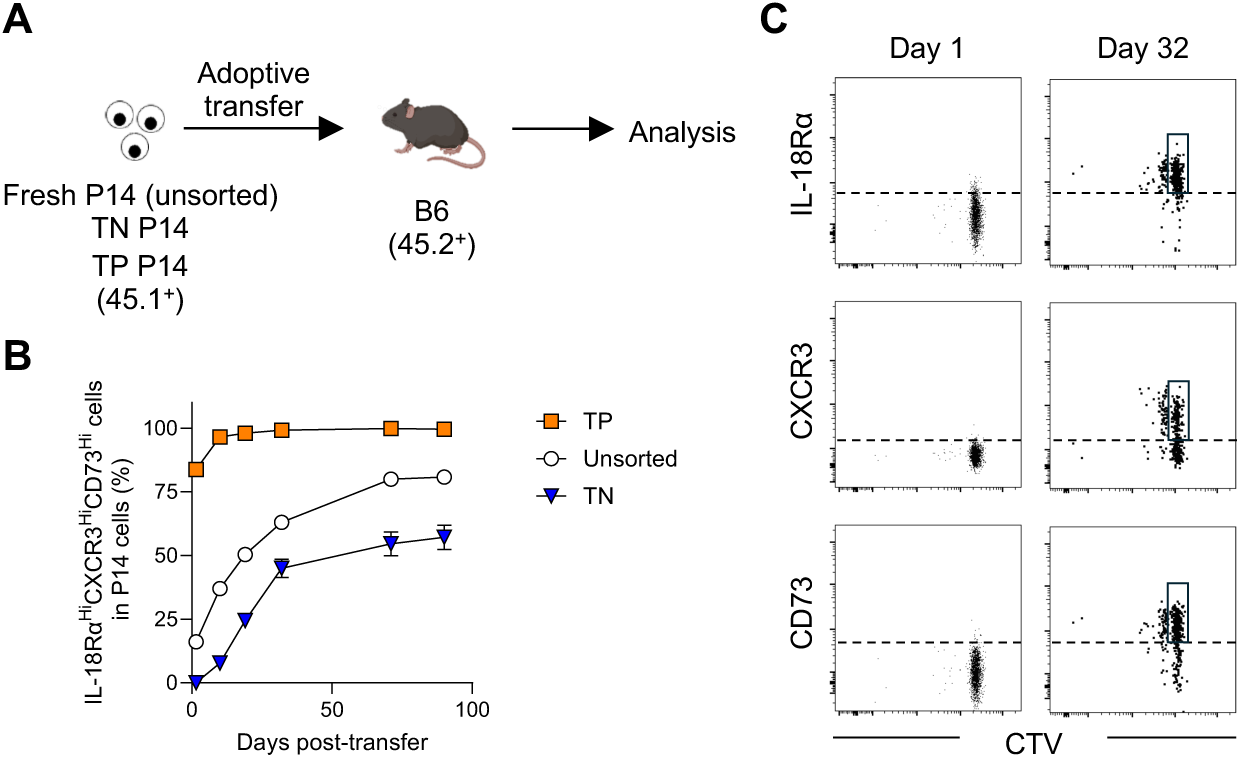
Naïve CD8 T cells alter their phenotype through differentiation. (A) Experimental design for panel L. Unsorted, triple positive (TP, IL-18Rα^Hi^CXCR3^Hi^CD73^Hi^), and triple negative (TN, IL-18Rα^Lo^CXCR3^Lo^CD73^Lo^) sorted P14 CD8 T cells were separately transferred into B6 mice. The phenotype of transferred P14 CD8 T cells was monitored in the blood over time. (B) Frequency of TP cells within transferred CD45.1^+^DbGP33^+^ P14 CD8 T cells. Data were pooled from 2 independent experiments with 3 or more mice per group. Each symbol represents the mean and error bars indicate SEM. (C) Sorted TN P14 CD8 T cells (CD45.1^+^) were stained with CTV (Celltrace violet) and adoptively transferred into recipient B6 mice (CD45.2^+^). The expression levels of IL-18Rα, CXCR3, and CD73 as well as cell division on transferred P14 CD8 T cells were analyzed on days 1 (PBMC) and 32 (spleen) post transfer. Representative flow cytometry plots were gated on DbGP33^+^ P14 CD8 T cells. Boxes highlight undivided marker^+^ cells.

### Naive CD8 T cell subsets differentially respond to infection

In Figure 2, we identified multiple naïve CD8 T cell subsets based on the expression of three markers and confirmed that the TP subset observed in long-lived P14 cells is also present in fresh P14 cells. However, whether the TP subset in fresh P14 cells possesses functional superiority similar to that of long-lived P14 cells remains unclear. Additionally, the response of other subsets to infection is yet to be determined. To address these questions, we sorted 5 subsets from fresh P14 cells: 1) IL-18Rα^Lo^CXCR3^Lo^CD73^Lo^ (TN subset), 2) IL-18Rα^Hi^CXCR3^Lo^CD73^Lo^, 3) IL-18Rα^Hi^CXCR3^Lo^CD73^Hi^, 4) IL-18Rα^Hi^CXCR3^Hi^CD73^Lo^, and 5) IL-18Rα^Hi^CXCR3^Hi^CD73^Hi^ (TP subset) (Fig. 5A). These sorted P14 cells were adoptively co-transferred with an equal number of congenically distinct fresh unsorted control P14 cells into B6 mice, followed by LCMV-Arm infection (Fig. 5A). Like long-lived P14 cells, the TP subset (subset 5) exhibited superior expansion when compared to unsorted control P14 cells (Fig. 5B). Interestingly, two double positive subsets, IL-18Rα^Hi^CXCR3^Lo^CD73^Hi^ (subset 3) and IL-18Rα^Hi^CXCR3^Hi^CD73^Lo^ (subset 4), also demonstrated enhanced responses relative to control P14 cells (Fig. 5B). In contrast, the single positive subset, IL-18Rα^Hi^CXCR3^Lo^CD73^Lo^ (subset 2), showed comparable kinetics to unsorted control cells, and the TN subset (subset 1) displayed significantly lower responses (Fig. 5B). To ensure that our observations were not limited to TCR transgenic CD8 T cells, we investigated responses of subsets within endogenous antigen-specific naïve CD8 T cells. IL-18Rα^Hi^CD73^Hi^ (double positive: DP) and IL-18Rα^Lo^CD73^Lo^ (double negative: DN) naïve CD8 T cells from B6 mice were sorted and separately transferred into congenically marked B6 mice, followed by LCMV-Arm infection (Fig. 5C and Fig. S5). Flow cytometric analysis revealed that donor IL-18Rα^Hi^CD73^Hi^ naïve DbGP33^+^ CD8 T cells expanded significantly more than IL-18Rα^Lo^CD73^Lo^ counterparts (Fig. 5, D and E). Collectively, these findings demonstrate that the phenotypic heterogeneity of naïve CD8 T cells correlate with their expansion capacity during infection.

**Fig. 5.**
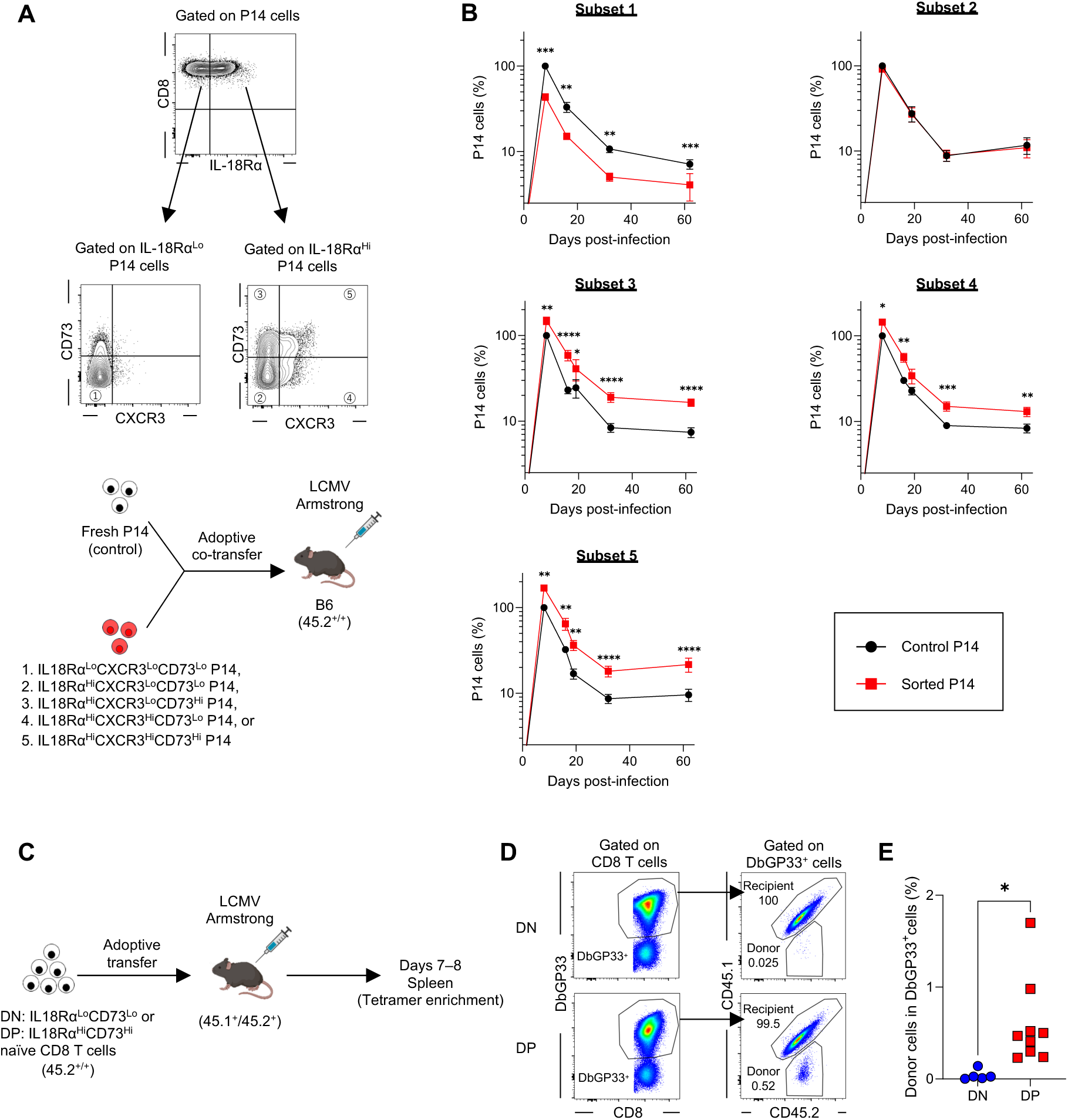
Naive CD8 T cell subsets differentially respond to infection. (A) Experimental design for panel B. Five naïve CD8 T cell subsets were sorted from the spleen of P14 TCR transgenic mice. Equal numbers (1 × 10^3^ cells each) of congenically distinct P14 CD8 T cells, including each sorted subset and freshly isolated P14 CD8 T cells (control), were adoptively co-transferred into B6 mice, followed by LCMV-Arm infection. The responses of sorted and control P14 populations in the blood were monitored over time. (B) Kinetics of the frequency of sorted and control P14 CD8 T cells. The frequency of control P14 cells at day 8 post-infection was set as 100. Each symbol represents the mean, and error bars indicate the SEM. (C) Experimental design for panels D and E. IL18Rα^Lo^CD73^Lo^ (DN: double negative) or IL18Rα^Hi^CD73^Hi^ (DP: double positive) CD44^Lo^ naïve CD8 T cells were sorted from splenocytes of B6 mice (CD45.2^+/+^). The number of DbGP33 tetramer^+^ CD8 T cells was confirmed using a portion of the sorted cells via tetramer enrichment. Bulk sorted cells, containing equal numbers of DbGP33 tetramer^+^ cells, were separately transferred into recipient mice (CD45.1^+^CD45.2^+^), followed by LCMV-Arm infection. On days 7–8 post-infection, tetramer enrichment was performed to analyze the response of donor-derived DbGP33^+^ CD8 T cells in the spleen. (D) Representative flow plots. (E) Frequency of donor-derived DbGP33^+^ CD8 T cells among total DbGP33^+^ CD8 T cells in the spleen. Each symbol represents an individual mouse. Data were pooled from 2 independent experiments with 5 or 9 mice per group. Statistical analyses were performed using one-way ANOVA (B) and unpaired *t* test (E). * *p* < 0.05; ** *p* < 0.01; ***: *p* < 0.001; ****: *p* < 0.0001.

We next tested whether the expansion capacity of naïve CD8 T cell subsets is linked to their protective abilities (West et al., 2011). Fresh P14, sorted TN P14, sorted TP P14, and long-lived P14 cells were separately transferred into B6 mice. On day 5 post-LCMV infection, spleen and liver were harvested from the recipient mice and the responses of P14 cells and viral titers were analyzed (Fig. 6A). Whereas viral titers in the TN group were slightly higher in the spleen than those in the fresh P14 group, the TP and long-lived P14 groups showed a substantial reduction in viral titers in the spleen and liver (Fig. 6, B and C). The expansion levels of P14 cells (Fig. 6D) were directly correlated with viral control. Consistent with Figures 1 and 5, the number of effector P14 cells in the spleen and liver was substantially higher in the TP and long-lived P14 groups compared to the fresh P14 group, even at this early time point (Fig. 6D). In contrast, the TN group exhibited a lower response (Fig. 6D). These data support the presence of functionally heterogenous naïve CD8 T cell populations with distinct protective abilities.

**Fig. 6.**
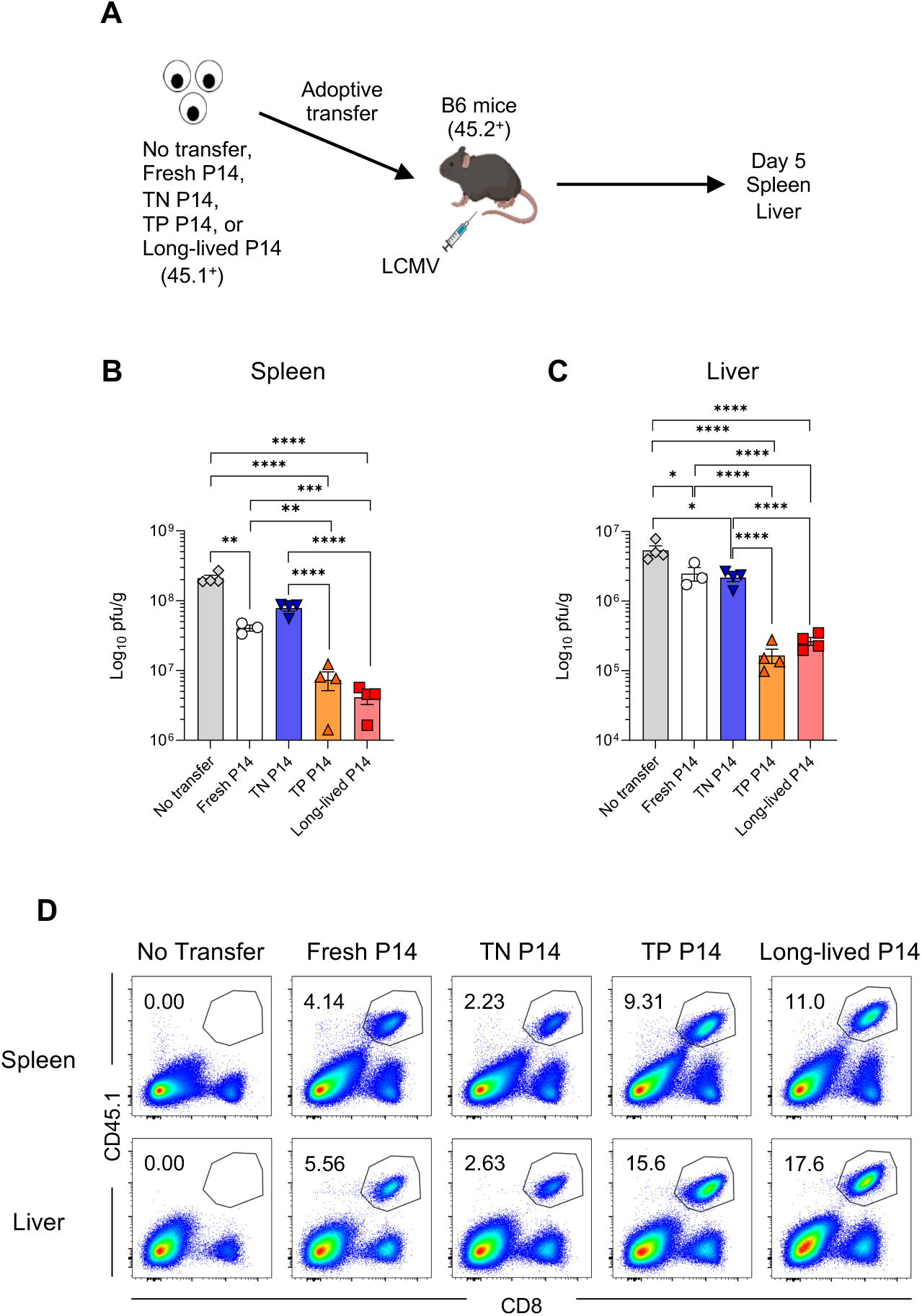
Naïve CD8 T cell subsets exhibit differential protective abilities. (A) Experimental design for panels B–D. Triple negative (TN, IL-18Rα^Lo^CXCR3^Lo^CD73^Lo^) and triple positive (TP, IL-18Rα^Hi^CXCR3^Hi^CD73^Hi^) P14 CD8 T cells were sorted from the spleens of P14 TCR transgenic mice. Freshly isolated P14 CD8 T cells, TN cells, TP cells, or long-lived P14 cells, housed for 34–50 days in B6 mice, (3 × 10^4^ cells/mouse, CD45.1^+^) were separately transferred into recipient B6 mice (CD45.2^+^). Mice were then infected with a low dose of LCMV-Clone 13 (2 × 10^4^ pfu/mouse). On day 5 post-infection, the spleen and liver were harvested, and viral titer was measured by plaque assay. (B and C) Quantification of viral titers in the spleen (B) and liver (C). (D) Representative flow cytometry plots, gated on live cells, show percentages of progeny derived from each naïve P14 CD8 T cell population (CD45.1^+^CD8 T cells) in the spleen and liver. Data are representative of 2 independent experiments with 3 or 4 mice per group. Each symbol represents an individual mouse, bars indicate the geometric mean, and error bars indicate SEM. Statistical analysis was performed using one-way ANOVA. * *p* < 0.05; ** *p* < 0.01; *** *p* < 0.001; **** *p* < 0.0001.

### Effector cells derived from long-lived naïve CD8 T cells exhibit enhanced survival during the T cell expansion phase

Our current study shows that naïve CD8 T cells comprise multiple subsets with distinct capacities to generate effector and memory cells. A critical question that needs to be answered is how functionally superior populations generate higher numbers of effector and memory CD8 T cells following infection. The increased number of virus-specific CD8 T cells after antigen stimulation could be explained by improved survival and/or accelerated proliferation. To examine these possibilities, long-lived P14 cells were adoptively co-transferred with an equal number of freshly isolated P14 cells into B6 mice. On day 5 post-LCMV-Arm infection, during the T cell expansion phase, the survival and proliferation of P14 cells were analyzed using annexin V/7-AAD and BrdU staining, respectively (Fig. 7A). Although both frequency and number of long-lived P14 cells were significantly higher than fresh P14 cells (Fig. 7, B and C), BrdU incorporation was similar between the two populations (Fig. 7, D and E), indicating no difference in proliferation. However, the frequency of live cells (Annexin V^Negative^7-AAD^Negative^ cells) was significantly higher in proliferating effector cells derived from long-lived P14 cells compared to those from fresh P14 cells at day 5 post-infection (Fig. 7, F and G). Taken together, these findings demonstrate that effector cells derived from long-lived P14 cells exhibit superior survival during the T cell expansion phase, contributing to the increased numbers of effector and memory CD8 T cells derived from this population.

**Fig. 7.**
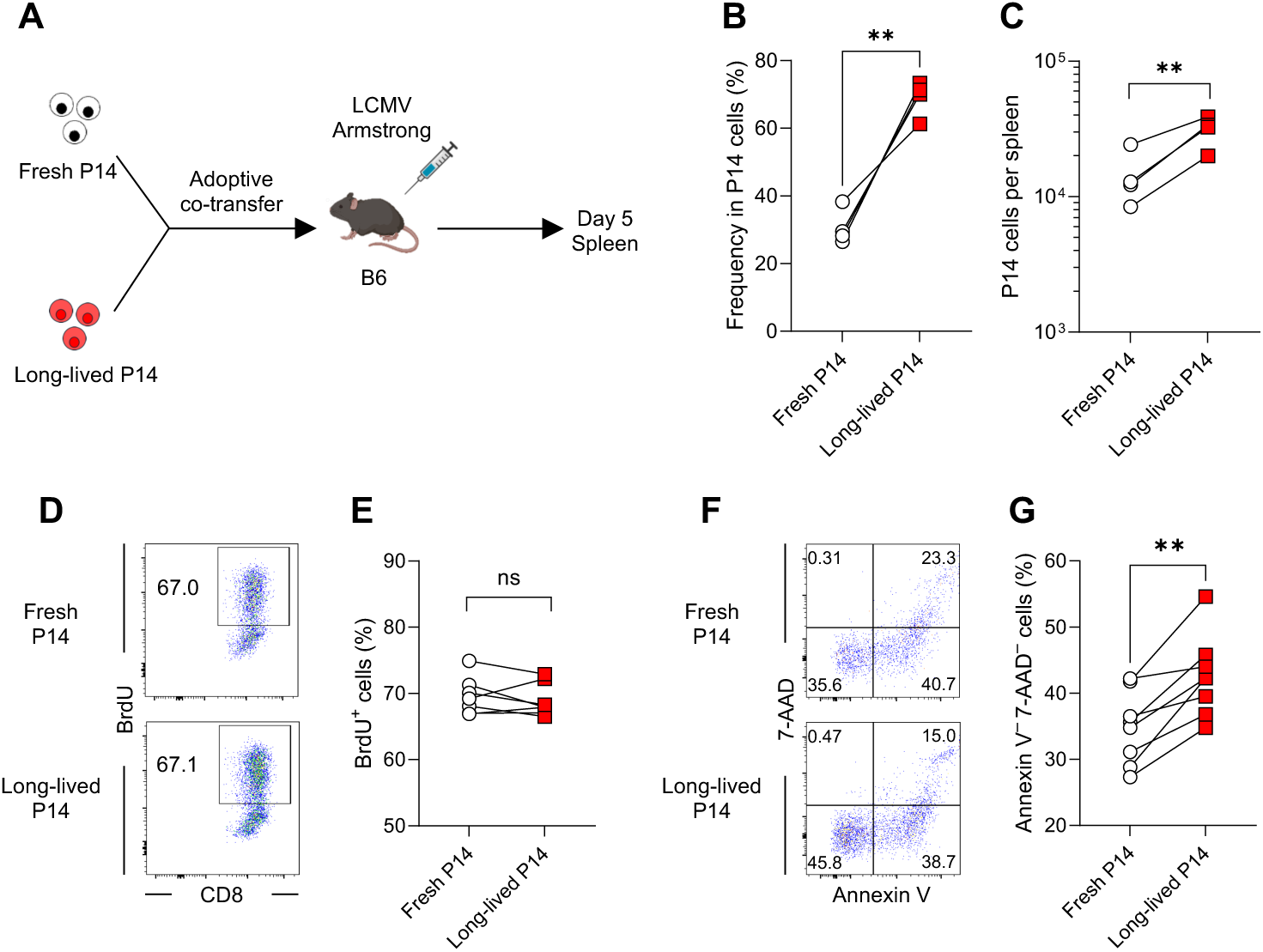
Effector cells derived from long-lived naïve CD8 T cells survive better during infection. (A) Experimental design for panels B–G. Congenically distinct long-lived (housed for 34 days in B6 mice) and freshly isolated P14 CD8 T cells (1 × 10^3^ cells each) were adoptively co-transferred into B6 mice, followed by LCMV-Arm infection. On day 5 post-infection, proliferation and survival of transferred P14 CD8 T cells in the spleen were assessed using a BrdU incorporation assay and annexin V/7-AAD staining, respectively. (B and C) Percentages of progeny derived from fresh and long-lived P14 CD8 T cells within total P14 CD8 T cells (B) and their absolute numbers (C). Data are representative of 4 independent experiments with 4 mice per group. (D–G) Frequency of BrdU^+^ P14 CD8 T cells (D and E) and annexin V^−^7-AAD^−^ live P14 CD8 T cells (F and G). Representative flow cytometry plots were gated on effector P14 CD8 T cells derived from either fresh- or long-lived-derived population. Data in panels E and G were pooled from 2 independent experiments with 7 or 8 mice per group. Each symbol represents an individual mouse, and lines indicate paired comparisons within the same mice. Statistical analysis was performed using paired *t* test. ns, not significant; ** *p* < 0.01.

### Ly6C2 is critical for enhanced survival of effector CD8 T cells during infection

To investigate the mechanisms underlying the superior survival of activated long-lived P14 cells post-infection, we compared the transcriptional profiles of effector CD8 T cells derived from long-lived and fresh P14 cells on day 5 post-infection. The RNA-seq analysis showed striking similarity between the two populations, with only one gene, *Ly6c2*, significantly upregulated in the progeny of long-lived P14 cells (Fig, 8A and Supplemental Table 2). This suggests a role for *Ly6c2* in the enhanced survival of effector CD8 T cells derived from long-lived P14 cells, leading us to focus on *Ly6c2* as a candidate molecule. First, we assessed Ly6C2 protein expression to confirm whether it was also upregulated in effector CD8 T cells derived from long-lived P14 cells. Due to the high homology between Ly6C2 and Ly6C1, available antibodies cannot distinguish these two molecules. However, since CD8 T cells predominantly express Ly6C2 (Morimoto et al., 2022), we used an anti-Ly6C antibody, which detects both molecules, to estimate Ly6C2 expression on CD8 T cells. Consistent with the RNA-seq data, Ly6C protein expression was significantly elevated in effector cells derived from long-lived P14 cells on day 5 post-infection (Fig. 8, B and C). Next, to evaluate the functional role of Ly6C2 upregulation in the enhanced survival of activated long-lived P14 cells post-infection, we generated long-lived P14 cells lacking Ly6C2 expression using CRISPR-Cas9-mediated knockout (KO) (Fig. S6).

**Fig. 8.**
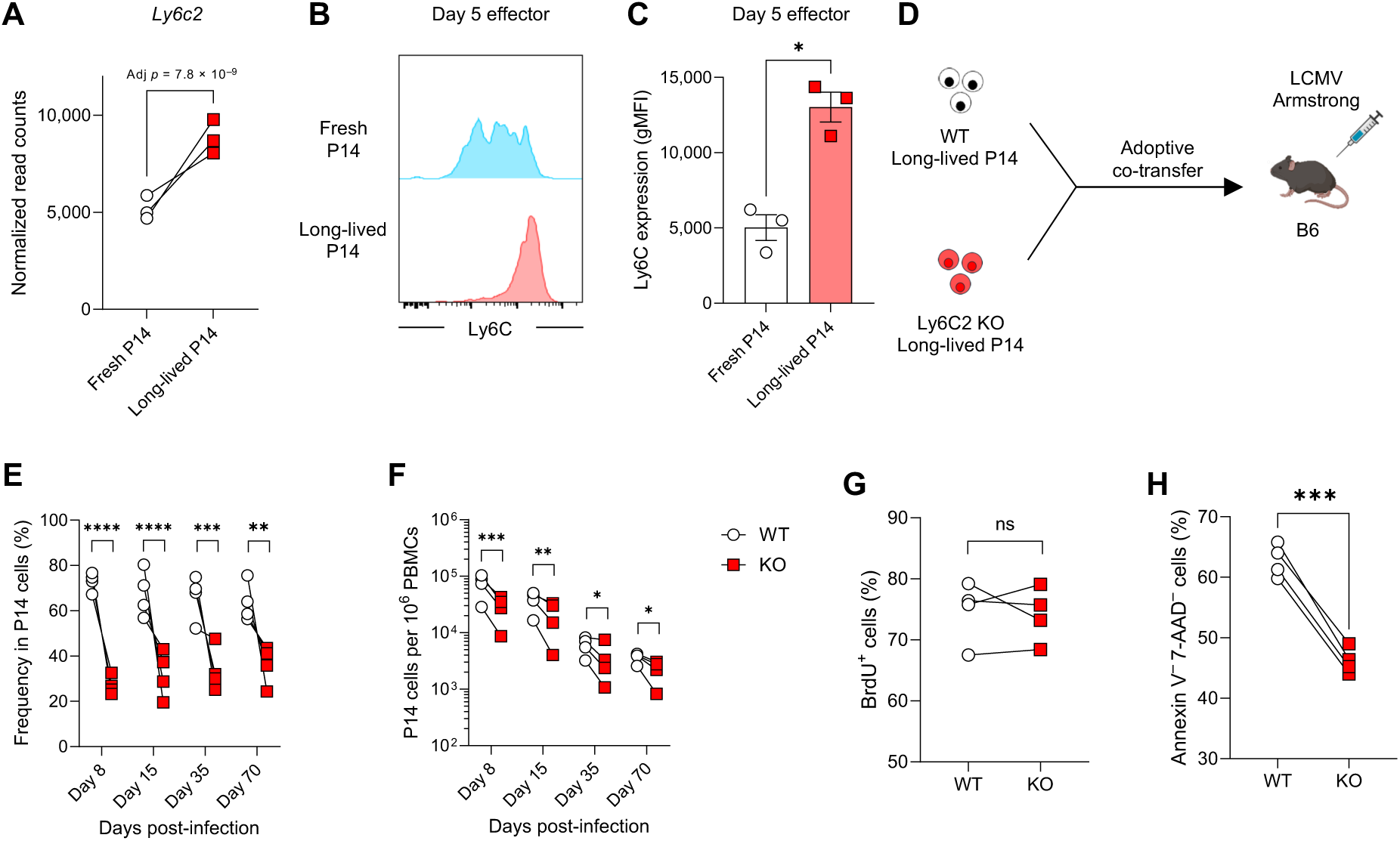
Ly6C2 on effector CD8 T cells plays a key role in enhancing their survival. (A) Normalized read counts for *Ly6c2* obtained from RNA-seq analysis are shown. Congenically distinct long-lived (housed for 57–65 days in B6 mice) and fresh P14 cells (1 × 10^3^ each) were adoptively co-transferred into B6 mice, followed by LCMV-Arm infection. On day 5 post-infection, effector P14 CD8 T cells derived from long-lived and fresh populations were sorted from the spleen for RNA-seq. (B and C) Ly6C protein expression on effector P14 CD8 T cells derived from fresh and long-lived P14 populations in the spleen on day 5 post-LCMV-Arm infection. Experimental design as in Figure 5A. Representative histograms were gated on P14 CD8 T cells derived from either fresh or long-lived P14 cells. Data are representative of 2 independent experiments with 3 mice per group. (D) Experimental design for panels E–H. Congenically distinct WT and Ly6C2 KO long-lived P14 CD8 T cells (1 × 10^3^ cells each) were adoptively co-transferred in B6 mice, followed by LCMV-Arm infection. Ly6C2 KO long-lived P14 CD8 T cells were generated using CRISPR-Cas9. (E and F) Percentages of progeny derived from WT and Ly6C2 KO long-lived P14 CD8 T cells within total P14 CD8 T cells (E) and their kinetics (F) in the blood post-infection. (G and H) Frequency of BrdU^+^ P14 CD8 T cells (G) and annexin V^−^7-AAD^−^ live P14 CD8 T cells (H) in the spleen on day 5 post-infection. Data are representative of 2 or 3 independent experiments with 4 mice per group. Each symbol represents an individual mouse. Statistical analyses were performed using Walt’s test (A), unpaired *t* test (C), paired *t* test (G and H), or one-way ANOVA (E and F). ns, not significant; * *p* < 0.05; ** *p* < 0.01; *** *p* < 0.001; **** *p* < 0.0001.

Consistent with a previous finding (Morimoto et al., 2022), our data confirm that CD8 T cells predominantly express Ly6C2 since anti-Ly6C antibody staining was substantially reduced following *Ly6c2* KO (Fig. S6). Ly6C2 KO long-lived P14 cells were adoptively co-transferred with an equal number of wild-type (WT) long-lived P14 cells into B6 mice, followed by LCMV-Arm infection (Fig. 8D). Compared to WT control P14 cells, the number of effector cells derived from Ly6C2 KO long-lived P14 cells was significantly reduced on day 8 post-infection, with this reduction persisting through the contraction and memory phases (Fig. 8, E and F). To determine if this reduction was due to impaired survival of Ly6C2 KO cells during the T cell expansion phase, as observed in Figures 7F and 7G, we performed annexin V/7-AAD and BrdU incorporation assays. While BrdU-incorporation was comparable between the two populations (Fig. 8G), the frequency of live effector cells derived from Ly6C2 KO long-lived P14 cells on day 5 post-infection was significantly lower than that of WT control cells (Fig. 8H). These data show that Ly6C2 plays a critical role in supporting the survival of activated long-lived P14 cells during infection.

## DISCUSSION

In this study, we have identified three cell surface markers (IL-18Rα, CXCR3, and CD73) that define functionally distinct naïve CD8 T cell subsets with hierarchical capacities to generate effector and memory cells. Specifically, the triple-negative subset exhibited the lowest potential, followed by the IL-18Rα single-positive subset. In contrast, subsets expressing IL-18Rα in combination with CXCR3, CD73, or both (triple-positive) generated significantly more effector and memory cells. This advantage was primarily driven by enhanced effector cell survival.

Mechanistically, Ly6C2 played a key role in promoting survival, as it was highly upregulated on effector cells derived from functionally superior naïve CD8 T cells, leading to their enhanced survival during T cell expansion phase. Furthermore, we observed that the triple-negative subsets could differentiate into the triple-positive subset, establishing a lineage relationship among these populations.

Several studies have attempted to define the heterogeneity of naïve CD8 T cells. One approach classified subsets based on tonic TCR signaling intensity, but findings have been contradictory: one study reported that naïve CD8 T cells receiving strong tonic TCR signals responded better to antigen stimulation, whereas another found the opposite (Eggert et al., 2024; Fulton et al., 2015). In our study, we observed that long-lived and freshly isolated naïve CD8 T cells similarly expressed CD5, suggesting that the subsets we identified are largely independent of tonic TCR signaling. Another attempt at elucidating naïve CD8 T cell heterogeneity is the study of recent thymic emigrants, the youngest naïve T cells (Fink, 2013). Previous studies have shown that these cells exhibit enhanced responses after infection compared to mature naïve T cells (Berkley and Fink, 2014; Deets et al., 2016). However, our findings cannot be attributed to recent thymic emigrants, as long-lived naïve CD8 T cells, which should contain fewer recent thymic emigrants, still demonstrated enhanced responses after infection. These observations suggest that functional changes of mature naïve CD8 T cells, rather than the presence of recent thymic emigrants, drive the differences in infection responses among the naïve CD8 T cell subsets identified in our study.

Previous studies identified Ly6C^Hi^ and Ly6C^Lo^ subsets within naïve CD8 T cells and reported that Ly6C^Hi^ naïve CD8 T cells generate more effector cells, although neither study determined whether Ly6C itself plays a functional role in CD8 T cell responses or merely serves as a marker (Berton et al., 2023; Ju et al., 2021). However, the findings regarding memory CD8 T cell formation were inconsistent. One study reported no difference in the final number of memory CD8 T cells derived from Ly6C^Hi^ versus Ly6C^Lo^ naïve cells (Ju et al., 2021), whereas the other found that Ly6C^Hi^ naïve CD8 T cells gave rise to a greater number of memory cells (Berton et al., 2023). A potential explanation for this discrepancy may lie in differences in type I IFN signaling. Ly6C expression in naïve CD8 T cells can be upregulated by type I IFN stimulation in the absence of cognate antigen, and such stimulation can also induce expression of effector-associated genes (Berton et al., 2023; Jergovic et al., 2021; Ju et al., 2021; Tu et al., 2025). Thus, the Ly6C^Hi^ naïve CD8 T cells in these studies may have been exposed to different levels or duration of type I IFN stimulation, potentially leading to these divergent outcomes in memory CD8 T cell formation. Our data further support that Ly6C expression alone is insufficient to define the functional potential of naïve CD8 T cells, since Ly6C^Lo^ long-lived naïve CD8 T cells responded to infection as effectively as the Ly6C^Hi^ subset. Collectively, these observations indicate that while Ly6C in naïve CD8 T cells can reflect prior exposure to type I IFN, it may not reliably indicate the memory-forming potential of naïve CD8 T cells. In contrast, our classification based on IL-18Rα, CD73, and CXCR3 expression identified naïve CD8 T cell subsets that gave rise to more memory precursor effector cells and ultimately produced a greater number of memory CD8 T cells. Therefore, this marker combination not only offers a more reliable predictor of memory-forming potential than Ly6C but also reveals previously unappreciated functional heterogeneity within the naïve pool. Mechanistically, although Ly6C expression in naïve CD8 T cells is not a critical indicator of functional potential, we found that *Ly6c2*—one of the two genes encoding Ly6C protein, distinct from *Ly6c1*—is selectively expressed in effector progeny derived from long-lived naïve CD8 T cells. Ly6C2 plays a critical role in promoting effector cell survival during the clonal expansion phase, thereby contributing to robust immune responses. This distinction underscores the context-dependent role of Ly6C: while limited as a predictive marker in the naïve state, it serves a functional role in supporting effector cell survival.

Our findings also suggest that naïve CD8 T cells could be targeted to enhance protective T cell immunity. Given that the naïve CD8 T cell population includes functionally distinct subsets, increasing the quantity of functionally superior subsets before stimulation could improve CD8 T cell responses to vaccines and infection. Although strategies to selectively expand these subsets remain to be determined, understanding how functionally inferior triple-negative naïve CD8 T cells differentiate into superior subsets will be key to developing approaches that enhance protective immunity.

Furthermore, our findings may provide a basis for a novel strategy to regulate CD8 T cell responses. We identified Ly6C2 as a critical molecule that promotes effector CD8 T cell survival. Consequently, blocking Ly6C2 could mitigate effector T cell responses by accelerating their death. Since CD8 T cell-mediated immunopathology can occur during viral infections and immune checkpoint blockade therapy for cancer (Iannacone and Guidotti, 2022; Ramos-Casals et al., 2020; Schmidt and Varga, 2018), inhibiting Ly6C2 may help improve patient outcomes in conditions with excessive CD8 T cell activity.

Despite several past studies (Berton et al., 2023; Eggert et al., 2024; Fulton et al., 2015; Jergovic et al., 2021; Ju et al., 2021), our understanding of naïve CD8 T cell heterogeneity remains limited compared to that of effector, memory, and exhausted CD8 T cells. Our findings provide evidence that naïve CD8 T cells are not a functionally uniform population but instead consist of distinct subsets with differential capacities to generate effector and memory cells. These results highlight a previously unrecognized layer of immune regulation that influences CD8 T cell responses. Future studies will be essential to elucidate the molecular mechanisms driving the differentiation of these subsets and to explore potential therapeutic strategies that leverage naïve CD8 T cell heterogeneity to improve immune protection against infections and cancer, and to regulate excessive CD8 T cell immunity.

## MATERIALS AND METHODS

### Mice

C57BL/6J (B6) mice (strain #: 000664) and CD45.1 B6 mice (strain #: 033076) were purchased from The Jackson Laboratory. To generate CD45.1^+^CD45.2^+^ B6 mice, female B6 mice were bred with male CD45.1 B6 mice. TCR transgenic P14 mice were bred and maintained in-house. All mice were used at 6-12 weeks of age and housed under specific pathogen-free conditions. All experiments were performed under the Institutional Animal Care and Use Committee of Cincinnati Children’s Hospital approved protocols (protocol # 2023-1039).

### Generation of long-lived naïve P14 cells

Naive P14 CD8 T cells (3–7 × 10^6^ cells/mouse, CD45.1^+^) were intravenously transferred into B6 mice (CD45.2^+^). The transferred P14 CD8 T cells were housed in the recipient B6 mice for more than 30 days. After *in vivo* housing, the housed P14 CD8 T cells were isolated from spleens and used for the experiments.

### Adoptive transfer and infection

For adoptive transfer experiments, a long-lived or sorted P14 CD8 T cell population (1 × 10^3^ cells/mouse) was adoptively co-transferred with an equal number of freshly isolated or short- housed P14 CD8 T cells (1 × 10^3^ cells/mouse) into recipient B6 mice. On the following day, the recipient B6 mice were infected with LCMV Armstrong (LCMV-Arm, 2 × 10^5^ pfu/mouse) by intraperitoneal injection, LCMV-Clone 13 (2 × 10^6^ pfu/mouse) by intravenous injection, or GP33 epitope-expressing *Listeria monocytogenes* (Shen et al., 1998; Shen et al., 2003) (3 × 10^3^ cfu/mouse) by intravenous injection. For the analysis of naïve P14 CD8 T cell phenotype over time, unsorted, sorted IL-18Rα^Hi^CD73^Hi^CXCR3^Hi^, or sorted IL-18Rα^Lo^CD73^Lo^CXCR3^Lo^ P14 cells (1–3 × 10^6^ cells/mouse, CD45.1^+^) were separately transferred into B6 mice (CD45.2^+^). In Figure 4C, CellTrace Violet Cell Proliferation Kit (Thermo Fisher Scientific) was used to label the sorted cells with celltrace violet prior to adoptive transfer. For the analysis of endogenous IL- 18Rα^Hi^CD73^Hi^ and IL-18Rα^Lo^CD73^Lo^ naïve CD8 T cell responses, IL-18Rα^Hi^CD73^Hi^CD44^Lo^ and IL-18Rα^Lo^CD73^Lo^CD44^Lo^ CD8 T cells were sorted from the spleen of naïve B6 mice using FACSAria Fusion (BD Biosciences) or FACSymphony S6 (BD Biosciences). To prevent anti- CD44 antibody-mediated depletion of sorted cells after adoptive transfer, recombinant anti- CD44 antibody (clone: REA664, Miltenyi Biotec), that has a mutated Fc region, was used.

Sorted cells (4.5 × 10^5^ cells/mouse, CD45.2^+/+^) were separately transferred into recipient B6 mice (CD45.1^+^/CD45.2^+^). On the following day, the recipient B6 mice were infected with LCMV-Arm as described above.

### Flow cytometry

Cells were isolated from the spleen, liver, LNs, and blood and incubated with an antibody cocktail for 30 minutes on ice. A LIVE/DEAD Fixable Near-IR Dead Cell Stain Kit (Invitrogen) was used to gate out dead cells. After surface staining, cells were fixed by using a Cytofix/Cytoperm kit (BD Biosciences). The following antibodies were purchased from BioLegend: CD8 (clone: 53-6.7), CD45.2 (clone: 104), CD45.1 (clone: A20), CD5 (clone: 53- 7.3), IL-18Rα (clone: A17071D), CXCR3 (clone: CXCR3-173), KLRG1 (clone: 2F1/KLRG1), CD127 (clone: A7R34), Ly6C (clone: HK1.4), and CD44 (clone: IM7). The following antibodies were purchased from Invitrogen: CD73 (clone: eBioTY/11.8) and CD62L (clone: MEL-14). The following antibodies were purchased from Miltenyi Biotec: CD44 (clone: REA664) and Ly6C (clone: REA796). APC-conjugated or BV421-conjugated DbGP33 MHC I tetramer was used for the detection of GP33-specific CD8 T cells. For annexin V and 7-AAD staining (Vezys et al., 2011), FITC Annexin V Apoptosis Detection Kit with 7-AAD (BioLegend) was used according to the manufacturer’s instructions. For the detection of BrdU incorporation, BrdU (1 mg/mouse) was intraperitoneally administered in mice (Araki et al., 2017). Two hours after injection, BrdU incorporation in splenic P14 cells was analyzed using FITC BrdU Flow Kit (BD Biosciences) following the manufacturer’s protocols. Samples were acquired on LSRFortessa (BD Biosciences) or FACSCanto (BD Biosciences). All FACS data were analyzed using FlowJo software (BD Biosciences).

### Protective capacity of naïve CD8 T cell subsets

Protective capacity of naïve CD8 T cells was examined as described previously (West et al., 2011). Briefly, freshly isolated P14, long-lived P14, sorted IL-18Rα^Hi^CD73^Hi^CXCR3^Hi^ P14 cells, or sorted IL-18Rα^Lo^CD73^Lo^CXCR3^Lo^ P14 cells (3 × 10^4^ cells/mouse, CD45.1^+^) were separately transferred into B6 mice (CD45.2^+^). On the following day, the recipient B6 mice were intravenously infected with LCMV-Clone 13 (2 × 10^4^ pfu/mouse). The spleen and liver obtained on day 5 post-infection were homogenized in Pre-Filled Bead Mill Tubes (Fisher Scientific) using Bead Ruptor Elite (OMNI international). LCMV titers in these homogenates were quantitated by plaque assay. Vero E6 cells (4 × 10^6^ cells/plate) were plated in 6-well plates (Corning), and the plates were incubated at 37°C under 5% CO_2_. On the following day, the plates (cell confluency: >90%) were used for the assay. The medium was removed from the plates, and 200 μL of titrated homogenate solution was added to the cells. After adsorption for an hour at 37°C under 5% CO_2_, the cells were overlaid with 4 mL of 0.5% Seakem ME Agarose (Lonza) in Medium 199 (Gibco) supplemented with 10% fetal bovine serum (FBS, Gibco), 1× Antibiotic-Antimycotic (Gibco), and 1× L-glutamine (Gibco). The plates were incubated for 4 days at 37°C under 5% CO_2_ and then overlaid with 2 mL of 0.5% agarose in Medium 199 containing 0.02% neutral red (Fisher Scientific). On the following day, plaques were counted.

### Tetramer enrichment

Fc blocking was performed by incubating splenic and LN single cell suspensions with anti-mouse CD16/32 antibody (clone: 93, BioLegend) for 15 minutes on ice. Cells were then stained with APC-conjugated and BV421-conjugated DbGP33 tetramers for 30 minutes on ice. After washing twice with MACS buffer which is PBS containing 2% FBS and 2 mM EDTA (invitrogen), cells were incubated with anti-APC microbeads (Miltenyi Biotec) for 15 minutes at 4°C. Cells were then washed, resuspended in MACS buffer, and the tetramer positive cells were enriched by using LS-columns (Miltenyi Biotec) according to the manufacturer’s instructions.

For Figure 5D, a portion of the sorted cells were used for tetramer enrichment to confirm that the sorted populations contain equal numbers of DbGP33 tetramer^+^ naïve CD8 T cells.

### Parabiosis

All surgeries were performed at the Veterinary Services Facility, Cincinnati Children’s Hospital Medical Center. Age-matched congenically marked B6 mice (CD45.1^+/+^ and CD45.2^+/+^) were conjoined by parabiosis surgery as described previously (Kamran et al., 2013). In brief, mice were anesthetized, surgically prepped by placing side by side in supine position, and administered pre-emptive analgesics. Longitudinal skin incisions were performed extending from the elbow to the knee, on the left side of one mouse and the right side of the other. The skin was gently dissected free from the underlying subcutaneous fascia. The surgeon joined the left olecranon of one animal to the right olecranon of the other using non-absorbable nylon suture, following the same procedure for the left and right knee joints. The surgeon then connected the skin of the two animals with absorbable suture. Wound clips and tissue adhesive were used to reinforce the suture line in areas of high tension surrounding the joints. Analgesia was continued for a minimum of 72-hours post-operatively. Wound clips were removed 14 days after surgery. The parabiosed mice were allowed to rest for 35 days. To reverse parabiosis, mice were anesthetized and prepped as described above. A longitudinal incision of the joined skin was made starting at the knee joint up to the elbow, freeing the skin from the subcutaneous fascia.

The sutures affixing the olecranon and knee joints were removed. The skin of each mouse was closed using absorbable suture. Analgesia was continued for a minimum of 72-hours post- operatively. 1.5–2.5 months after reversal surgery, the phenotype of host- and donor-derived bulk CD44^Lo^ and DbGP33^+^CD44^Lo^ CD8 T cells was analyzed using tetramer enrichment as described above.

### Bulk RNA-sequencing analysis

To analyze transcriptional differences between fresh and long-lived P14 cells prior to infection, P14 CD8 T cells (5 × 10^6^ cells/mouse) were transferred into B6 mice, and transferred P14 cells were re-isolated on day 1.5 or 35 post-transfer for cell sorting. To eliminate possible environmental differences, short-housed P14 cells were used for this analysis instead of freshly isolated P14 cells. RNA was extracted from these sorted cells using Trizol reagent (Life Technologies) and RNA Clean and Concentrator-5 (Zymo Research). The Bulk RNA-Seq library was made using Illumina Standard Total RNA Prep with Ribo-Zero plus kit (Illumina) and sequencing analysis was performed by Novogene Co. RNA-Seq data are available at the GEO database (accession GSE290461). To analyze transcriptional differences between effector P14 cells derived from fresh and long-lived P14 cells, freshly isolated and long-lived P14 cells were adoptively co-transferred into recipient B6 mice, followed by LCMV-Arm infection as described above. On day 5 post-infection, spleens were harvested from the recipient mice and fresh P14- and long-lived P14-derived populations were sorted. RNA was extracted as described above.

Bulk RNA-Seq library was made by the Single Cell Genomics Facility, Cincinnati Children’s Hospital Medical Center, using Illumina Nextera XT DNA Library Prep kit (Illumina) and sequencing analysis was performed by the Genomics Sequencing Facility, Cincinnati Children’s Hospital Medical Center. RNA-Seq data are available at the GEO database (accession GSE290460). Data from the bulk RNA-seq analyses were analyzed using Galaxy (Afgan et al., 2018) and R as described previously (Ando et al., 2023). The Poisson distance between samples was calculated using the PoiClaClu R package.

### CRISPR/Cas9 based Ly6C2 knockout

For gene knockout experiments, a sgRNA targeting the murine *Ly6c2* (5′- ACUGUGUGCAGGAAGAGGUG-3′) was purchased from Synthego. CD8 T cells were purified from the spleen of B6 mice that received P14 CD8 T cells >30 days prior, using the mouse CD8a^+^ T Cell Isolation Kit (Miltenyi Biotec) according to the manufacturer’s instructions. To generate Ly6C2 KO CD8 T cells using a CRISPR/Cas9 based gene editing system, purified CD8 T cells were electroporated with 6 μg of Alt-R S.p. Cas9 Nuclease (IDT) and 0.3 nmol of Ly6C2-targeted sgRNA using Lonza 4D-Nucleofector (Lonza) as described previously (Nussing et al., 2020). As a WT control, isolated CD8 T cells were electroporated only with Cas9 Nuclease. For P14 experiments, congenially marked Ly6C2 KO and WT long-lived P14 cells (1 × 10^3^ cells each) were adoptively co-transferred into B6 mice, and the recipient mice were immediately infected with LCMV-Arm.

### Statistical analysis

Statistical analyses except for RNA-seq data were performed with Prism 10 (GraphPad). For two-group comparisons, *p* values were determined by 2-tailed paired *t* test or 2-tailed unpaired *t* test. For multi-group comparison, *p* values were determined by one-way ANOVA with Šidák’s or Tukey’s multiple comparisons test. Log-transformed values were used for statistical comparisons in figures with a log scale. Statistical analyses for RNA-seq data were performed with Deseq2. Adjusted *p* values were determined by Walt’s test.

## Supplemental Materials

Figure S1 shows the phenotype of effector CD8 T cells derived from fresh and long-lived P14 cells on Day 8 post-LCMV-Arm infection, as well as the frequency and number of memory CD8 T cells derived from the two populations in the spleen on day 98 post-infection. Figure S2 shows the responses of fresh and long-lived P14 cells during *Listeria monocytogenes* infection. Figure S3 shows CD5 and Ly6C expression in fresh and long-lived P14 cells, along with the responses of Ly6C^Lo^ and Ly6C^Hi^ long-lived P14 cells during LCMV-Arm infection. Figure S4 shows the expression levels of IL-18Rα, CXCR3, and CD73 in fresh and long-lived P14 cells isolated from the spleen, liver, LNs, and blood. Figure S5 shows the cell sorting strategy for Figure 5C. Figure S6 shows the efficiency of Ly6C2 KO in long-lived P14 CD8 T cells. Supplemental Table 1 provides row data from transcriptional analysis of fresh and long-lived naïve P14 CD8 T cells prior to infection. Supplemental Table 2 provides row data from transcriptional analysis of effector cells derived from fresh and long-lived naïve P14 CD8 T cells on day 5 post-infection.

## Acknowledgments

We would like to thank the Veterinary Services Facility and Research Flow Cytometry Facility at Cincinnati Children’s Hospital Medical Center. We also would like to acknowledge the Single Cell Genomics Facility and Genomics Sequencing Facility at Cincinnati Children’s Hospital Medical Center for their assistance with bulk RNA-seq library preparation and sequencing, respectively. We would like to thank Drs. Chandrashekhar Pasare (Cincinnati Children’s Hospital Medical Center) and Sing Sing Way (Cincinnati Children’s Hospital Medical Center) for discussion and critical reading of this manuscript. Figures for experimental design were created using BioRender (BioRender.com).

This work was supported by Startup funds from the Cincinnati Children’s Hospital Research Foundation (to KA) and NIH grants R01AI139675 (to KA), R01AI184466 (to KA), and T32AI165396 (to YS). YS was supported by Japan Society for the Promotion of Science Overseas Research Fellowships.

## Author contributions

Yamato Sajiki: Conceptualization, investigation, data curation, formal analysis, funding acquisition, visualization, validation, and writing—original draft, review, and editing. Satomi Ando: Conceptualization, investigation, formal analysis, and writing—review and editing. Charles M. Perkins: Investigation, and writing—review and editing. Yi-Chung Huang: Investigation, and writing—review and editing. Katie E. Smith: Investigation, methodology, and writing—review and editing. Carolyn M. Doerning: Investigation, methodology, and writing—review and editing. Koichi Araki: Conceptualization, investigation, funding acquisition, visualization, validation, project administration, supervision, and writing— original draft, review, and editing.

## Data and materials availability

All data generated in the present study are present in the manuscript or supplemental materials and will be shared by the corresponding author upon reasonable request. Raw data files of the RNA-seq analyses on fresh and long-lived naïve P14 cells prior to infection and effector P14 cells derived from fresh and long-lived cells on day 5 post-infection have been deposited in the NCBI GEO database under the accession numbers, GSE290461 and GSE290460, respectively.

---

**Fig. S1.**
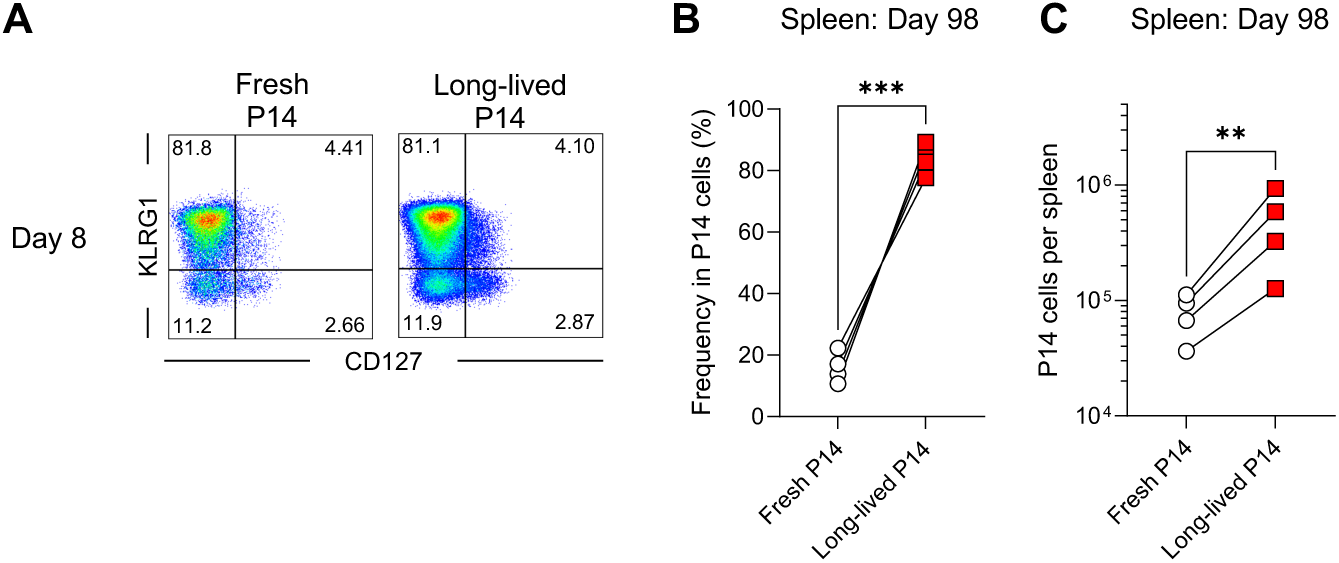
Response and differentiation of long-lived naive CD8 T cells after infection. Experimental design is shown in Figure 1C. (A) The phenotype (KLRG1 and CD127 expression) of progeny derived from fresh and long-lived P14 CD8 T cells in the blood on day 8 post-LCMV-Arm infection. Representative flow cytometry plots were gated on P14 CD8 T cells derived from either fresh- or long-lived-derived population. (B and C) Percentages of progeny derived from fresh and long-live P14 CD8 T cells within total P14 CD8 T cells (B) and their absolute numbers (C) in the spleen at day 98 post-LCMV-Arm infection. Results were representative of 3 or more independent experiments with 4 mice per group. In B and C, each symbol represents an individual mouse, and lines indicate paired comparisons within the same mice. Statistical analysis was performed using paired *t* test. ** *p* < 0.01; *** *p* < 0.001.

**Fig. S2.**
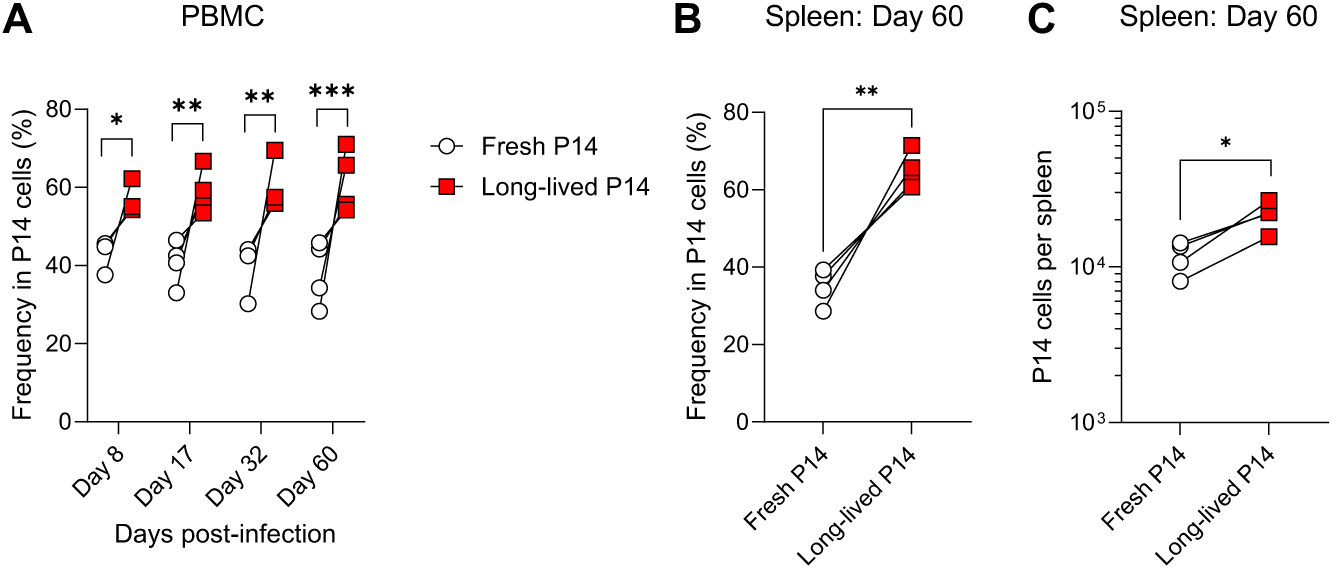
Responses of long-lived naïve CD8 T cells during *Listeria monocytogenes* infection. (A–C) Responses of long-lived P14 CD8 T cells during *Listeria monocytogenes* infection. Long-lived P14 CD8 T cells (1 × 10^3^ cells, CD45.1^+/+^), housed for 40 days in B6 mice, were adoptively co-transferred with an equal number of fresh P14 cells (1 × 10^3^ cells, CD45.1^+^CD45.2^+^) into B6 mice (CD45.2^+/+^), followed by GP33 epitope-expressing *Listeria monocytogenes* infection. Percentages of progeny derived from each population within total P14 CD8 T cells in the blood (A) and Spleen (B). (C) The absolute numbers of memory cells derived from fresh and long-lived P14 cells in the spleen at day 60 post-infection. Data are representative of 2 independent experiments with 4 mice per group. Each symbol represents an individual mouse, and lines indicate paired comparisons within the same mice. Statistical analyses were performed using one-way ANOVA (A) and paired *t* test (B and C). * *p* < 0.05; ** *p* < 0.01; *** *p* < 0.001.

**Fig. S3.**
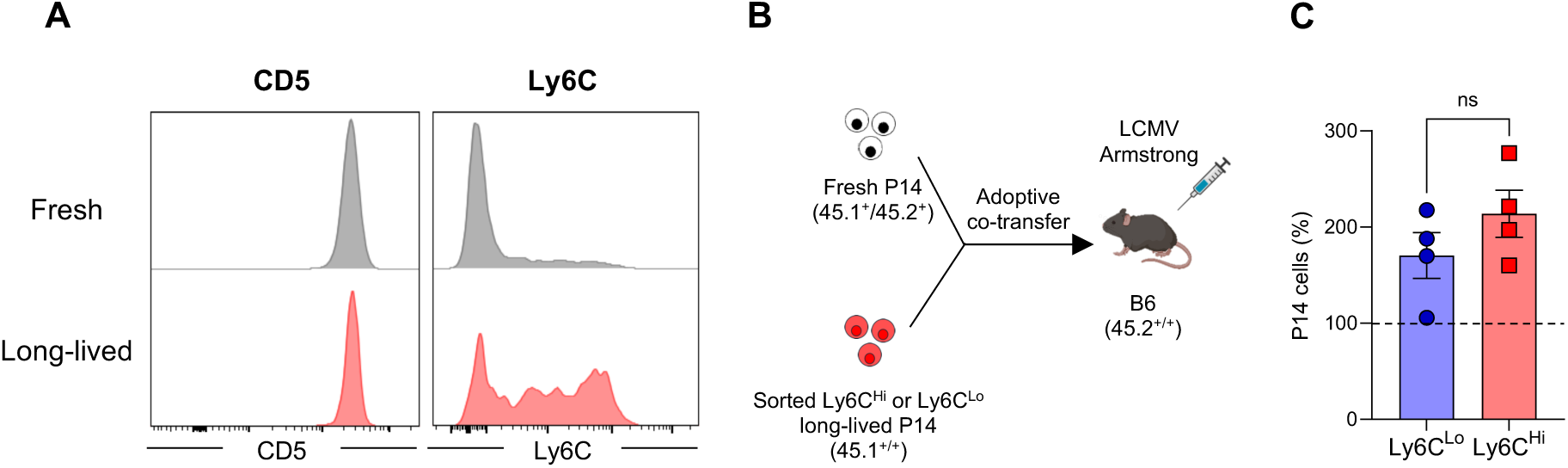
CD5 and Ly6C expression on fresh and long-lived P14 cells. (A) CD5 and Ly6C expression on long-lived (housed in B6 mice for more than 30 days) and fresh P14 CD8 T cells in the spleen. Representative histograms were gated on DbGP33^+^ P14 CD8 T cells. (B) Experimental design for panel C. Ly6C^Lo^ and Ly6C^Hi^ long-lived P14 CD8 T cells were sorted, and equal numbers (1 × 10^3^ cells each) of congenically distinct P14 CD8 T cells, including each sorted subset and freshly isolated P14 CD8 T cells, were adoptively co-transferred into B6 mice, followed by LCMV-Arm infection. The responses of sorted and control P14 populations in the blood were analyzed on day 8 post-infection. (C) Normalized frequency of sorted P14 CD8 T cells post-infection. The frequency of control P14 cells at day 8 post-infection was set as 100. Each symbol represents an individual mouse, bars indicate the mean, and error bars indicate SEM. Statistical analysis was performed using unpaired *t* test. ns, not significant.

**Fig. S4.**
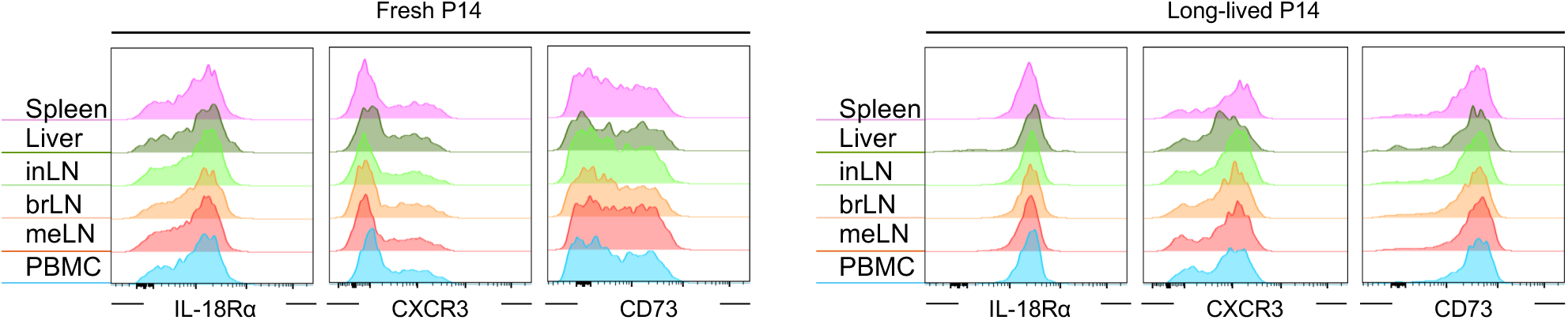
Heterogeneity of naïve CD8 T cells in different tissues. IL-18Rα, CXCR3, and CD73 expression on long-lived (housed in B6 mice for 39 days) and fresh P14 CD8 T cells in the spleen, liver, lymph nodes (LNs), and blood. Representative histograms were gated on DbGP33^+^ P14 CD8 T cells. inLN, inguinal LN; brLN, brachial LN; meLN, mesenteric LN.

**Fig. S5.**
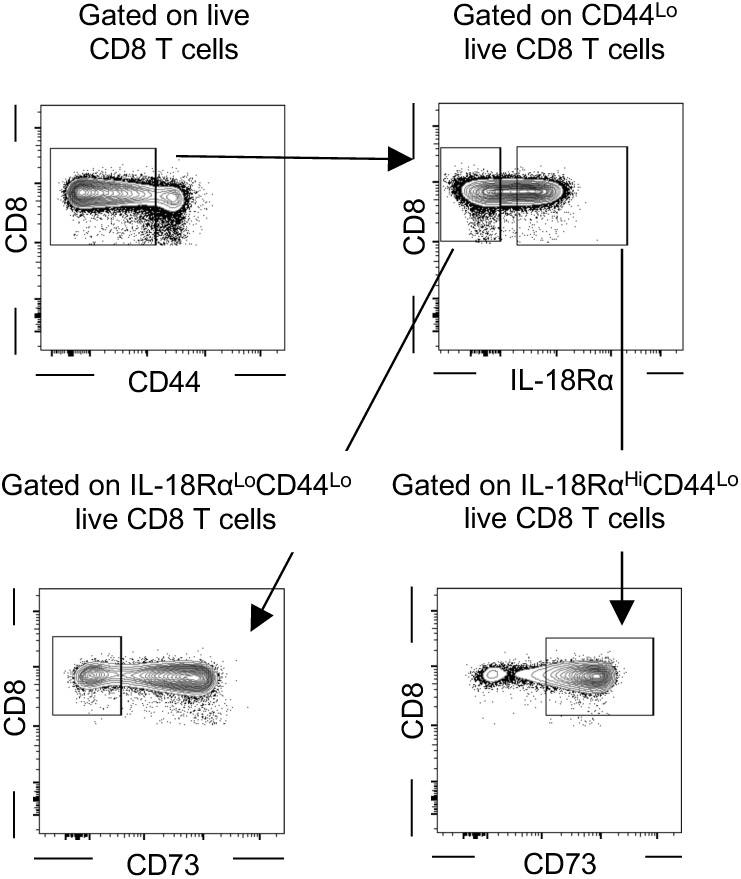
Cell sorting strategy for Figure 5C. IL-18Rα^Hi^CD73^Hi^CD44^Lo^ and IL-18Rα^Lo^CD73^Lo^CD44^Lo^ CD8 T cells were sorted from splenocytes of naïve B6 mice.

**Fig. S6.**
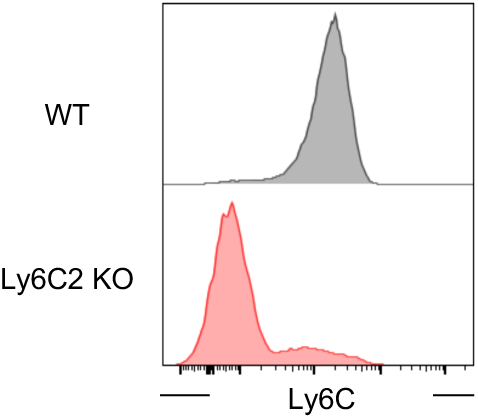
Efficiency of Ly6C2 KO in long-lived P14 CD8 T cells. Experimental design is shown in Figure 6D. Ly6C2 KO long-lived P14 CD8 T cells were generated using CRISPR-Cas9. Ly6C expression on WT and Ly6C2 KO long-lived P14 CD8 T cells in the blood on day 8 post-LCMV-Arm infection. Representative histograms were gated on effector P14 CD8 T cells derived from either WT or Ly6C2 KO long-lived P14 CD8 T cells.

